# Marker Data Enhancement For Markerless Motion Capture

**DOI:** 10.1101/2024.07.13.603382

**Authors:** Antoine Falisse, Scott D. Uhlrich, Akshay S. Chaudhari, Jennifer L. Hicks, Scott L. Delp

**Author notes:** This study was funded by the U.S. National Institutes of Health (NIH) under grant 1P41EB027060 and the Joe and Clara Tsai Foundation through the Wu Tsai Human Performance Alliance.

## Abstract

**Objective:** Human pose estimation models can measure movement from videos at a large scale and low cost; however, open-source pose estimation models typically detect only sparse keypoints, which leads to inaccurate joint kinematics. OpenCap, a freely available service for researchers to measure movement from videos, addresses this issue using a deep learning model— the marker enhancer—that transforms sparse keypoints into dense anatomical markers. However, OpenCap performs poorly on movements not included in the training data. Here, we create a much larger and more diverse training dataset and develop a more accurate and generalizable marker enhancer.

**Methods:** We compiled marker-based motion capture data from 1176 subjects and synthesized 1433 hours of keypoints and anatomical markers to train the marker enhancer. We evaluated its accuracy in computing kinematics using both benchmark movement videos and synthetic data representing unseen, diverse movements.

**Results:** The marker enhancer improved kinematic accuracy on benchmark movements (mean error: 4.1°, max: 8.7°) compared to using video keypoints (mean: 9.6°, max: 43.1°) and OpenCap’s original enhancer (mean: 5.3°, max: 11.5°). It also better generalized to unseen, diverse movements (mean: 4.1°, max: 6.7°) than OpenCap’s original enhancer (mean: 40.4°, max: 252.0°).

**Conclusion:** Our marker enhancer demonstrates both accuracy and generalizability across diverse movements.

**Significance:** We integrated the marker enhancer into OpenCap, thereby offering its thousands of users more accurate measurements across a broader range of movements.

## I. Introduction

MARKERLESS motion capture has become popular for biomechanical analysis of human movement because it reduces the cost and time associated with marker-based motion capture and facilitates large-scale, out-of-lab studies. Multi-camera video systems have achieved kinematic accuracy within approximately five degrees of marker-based motion capture for movements including walking, running, cycling, squatting, sit-to-stand, and drop jump [1]–[3]. These systems typically identify, for each video, the two-dimensional (2D) position of keypoints on the body using pose estimation models [4], reconstruct their three-dimensional (3D) position using triangulation algorithms like Direct Linear Transformation [5], and compute joint kinematics using, for example, multi-body models and inverse kinematics [6].

Several open-source pose estimation models can estimate 2D keypoints from video (e.g., OpenPose [7], HRNet [8]– [11], VITPose [12], and others [13]). These models, trained on datasets like COCO [14] and MPII [15] which have a limited number of body points labeled by non-experts in human anatomy, usually only detect a sparse set of joint centers [16]. This sparsity combined with the inherent noise of pose estimation models [17]–[19] make it difficult to estimate joint kinematics accurately. In previous work that compared a dual-camera system to marker-based motion capture [1], we found joint kinematic errors of up to 43° when using keypoints estimated from HRNet with a multi-body model and inverse kinematics. The large errors were primarily in the lumbar extension and hip flexion degrees of freedom, because of the few keypoints identified on the torso (shoulder joints), pelvis (hip joints), and femurs (knee joints). These keypoints fail to sufficiently constrain the kinematic redundancy of the multibody model, where various poses can produce identical joint center positions.

OpenCap is an open-source multi-camera system to compute the kinematics and kinetics of human movement from video [1]. To mitigate the problems that arise from keypoint sparsity, OpenCap incorporates a Long Short-Term Memory (LSTM) model—named the marker enhancer—that extrapolates the 3D position of 43 denser anatomical markers from the 3D position of 20 sparser keypoints (Fig. 1). We found that using the extrapolated anatomical markers reduced kinematic errors by 4.2° on average and by up to 35.9° at certain degrees of freedom compared to using video keypoints directly [1]. OpenCap, however, has lower performance on movements not included in the dataset used to train the marker enhancer (see examples in Fig. 2, column Uhlrich et al. (2023)). This is particularly noticeable in movements where the individual is lying prone, supine, or not executing movements aligned with the forward direction of the world frame, since the training set only includes movements performed upright and forward-facing. Ruescas-Nicolau et al. developed a comparable marker enhancer model using a different dataset and also observed varying performance across movements, with worse results on running compared to walking, squatting, and jumping [20].

**Fig. 1.**
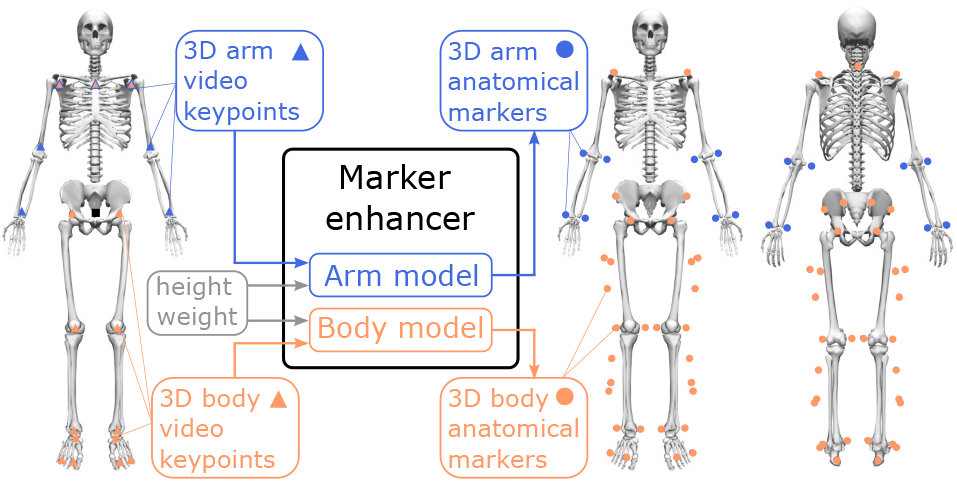
The marker enhancer model predicts the 3D position of 43 anatomical markers (colored dots on right skeletons) from 20 video keypoint (colored triangles on left skeleton) positions. It consists of two models: the *arm model* predicts the 3D position of eight arm-located anatomical markers from seven arm and shoulder keypoint positions (blue) and the *body model* predicts the 3D position of 35 anatomical markers located on the shoulder, torso, and lower-body from 15 shoulder and lower-body keypoint positions (orange). Both models include the subject’s height and weight as input, and all marker positions are expressed with respect to a root marker (the midpoint of the hip keypoints; black square on left skeleton).

**Fig. 2.**
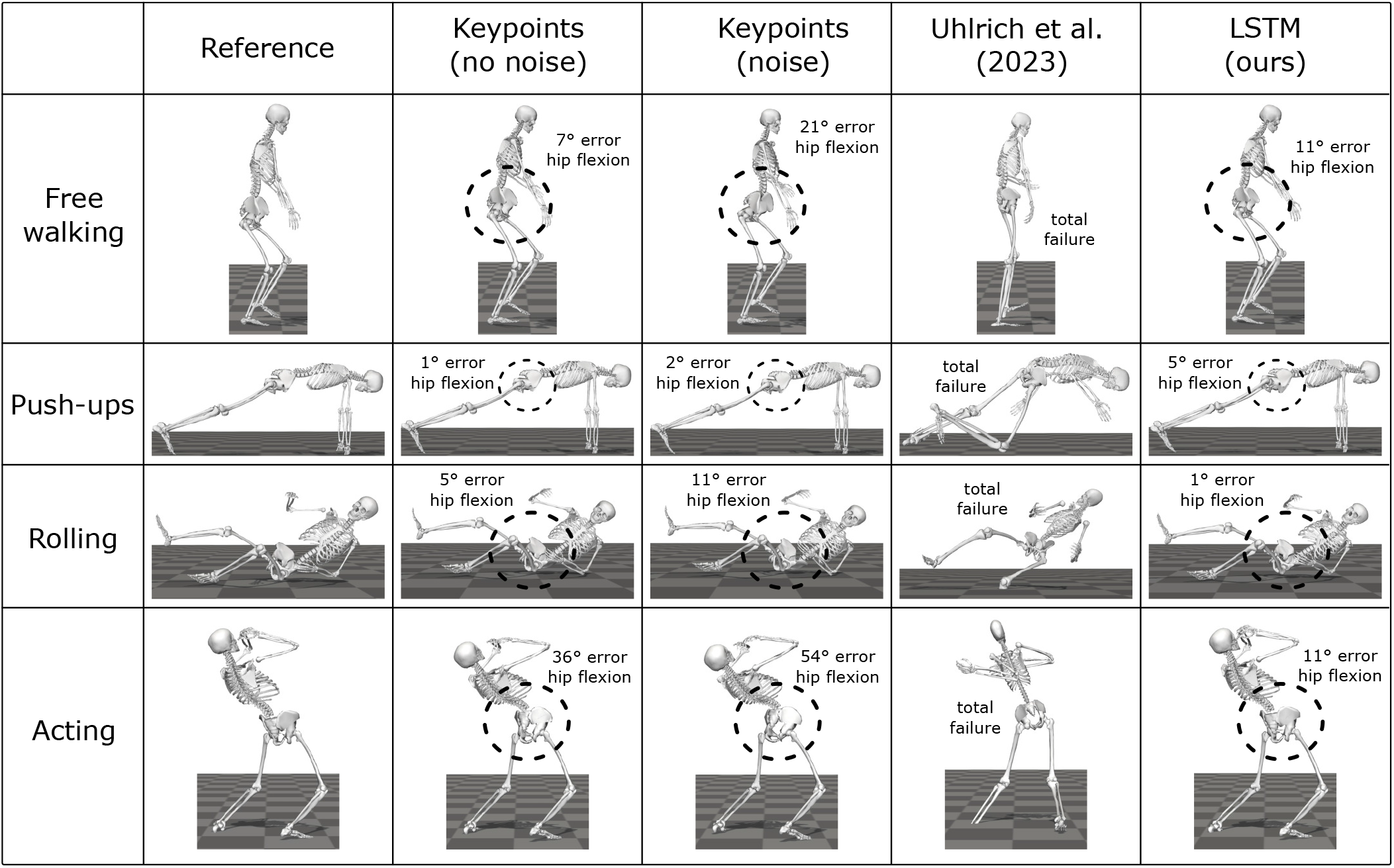
Kinematics of diverse movements computed from different synthetic marker sets: reference anatomical markers, video keypoints without and with noise, and anatomical markers predicted from noisy video keypoints using the marker enhancer from Uhlrich et al. (2023) [1] and our marker enhancer (LSTM) trained on the enhanced dataset. The instances shown are selected from the *generalizability* task dataset.

The objective of this study was to improve OpenCap’s ability to capture the kinematics of a broad range of movements (i.e., generalizability) while preserving its previously reported accuracy on a set of benchmark movements. We first created an enhanced dataset with over twice the data, or ten times more when including augmented data, from a much broader range of movements. We then trained three marker enhancers with different deep learning architectures: linear, LSTM, and transformer. We evaluated the performance of the three marker enhancers when computing joint kinematics from videos on a set of four benchmark movements. We then evaluated their ability to reconstruct joint kinematics from an unseen dataset containing a wide array of movements, thereby assessing their ability to generalize. Finally, we used muscle-driven tracking simulations to estimate dynamics from videos and evaluated the effect of more accurate joint kinematics on dynamic quantities. We integrated the best performing marker enhancer as part of the web-deployed version of OpenCap (https://www.opencap.ai/), and shared code and data to facilitate broader usage and reproduction of our work. A preliminary version of this work has been reported at [21].

## II. Methods

### A. Dataset

We compiled a large dataset of expert-processed marker-based motion capture data, and synthesized corresponding 3D video keypoints (n=20) and anatomical markers (n=43) to train the marker enhancers. We first processed the marker data from 16 movement datasets [22]–[39] with OpenSim [40] and AddBiomechanics [41] to obtain scaled OpenSim models and coordinate files (i.e., kinematic data). We then added virtual markers to the scaled models corresponding to the video keypoints and anatomical markers (Fig. 1). We finally extracted the 3D trajectory of each virtual marker for each coordinate file to create the synthetic dataset. To augment the experimental data, we simulated shorter and taller subjects by uniformly scaling each OpenSim model in the dataset by 90, 95, 105, and 110%, thereby quintupling the dataset size.

Prior to model training, we sampled the data at 60 Hz, split them into overlapping (50%) time-sequences of 1 s, and balanced the contribution of the individual datasets to obtain a wide movement distribution (Table I). In total, we included data from 1176 subjects and used over 221 hours of data (excluding data rotation, see next section). The dataset consisted of diverse movements, with the *Other* category, which includes a broad range of movements (e.g., dancing, cutting, stair climbing, squatting, jumping, rebounding in basketball, kicking and heading a soccer ball, performing hamstring exercises, balancing, sitting, boxing, and practicing yoga), accounting for 58% of the dataset (Table I). In comparison, OpenCap’s original marker enhancer had been trained on 108 hours of data from 336 subjects, from which over 80% was treadmill gait data and 45% was from a single dataset.

**Table I.**
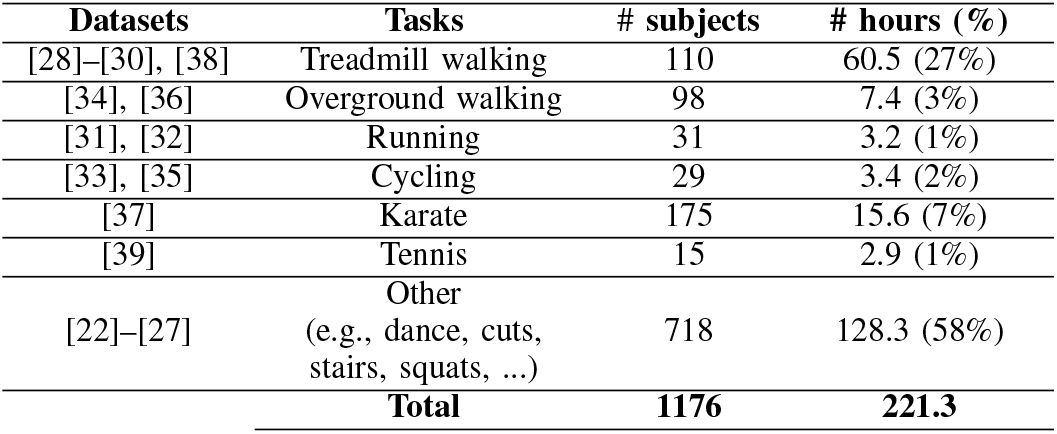
Dataset Distribution.

### B. Model architecture and training

We trained two models: one to predict anatomical markers located on the arms (*arm model*) and one to predict anatomical markers located on the shoulders, torso, and lower-body (*body model*) (Fig. 1). Since not all individual datasets included arms, we used a subset of the data (about 152 hours, i.e., 69% of total data) to train the *arm model*. The *body model* takes as input the 3D position of 15 shoulder and lower-body keypoints and predicts the 3D position of 35 anatomical markers, whereas the *arm model* uses the position of seven arm and shoulder keypoints to predict the position of eight anatomical markers. Both models include the subject’s height and weight as input.

During training, we expressed the 3D position of each marker with respect to a root marker (the midpoint of the hip keypoints), normalized the root-centered 3D positions by the subject’s height, added Gaussian noise (standard deviation: 18 mm) to each time step of the normalized positions [17]– [19], and standardized the data to have zero mean and unit standard deviation. Finally, to represent movements performed in different directions, we rotated each sample eight times; we applied six rotations about the vertical axis, and two random Euler rotation sequences to represent movements performed in any orientation. In total, when including rotations (i.e., data augmentation), we used 1433 hours of data for training the model, compared to the 85 hours used in the training set of the original OpenCap model.

We compared three deep learning architectures for our marker enhancer: linear, LSTM, and transformer models. For the LSTM models, we performed a random search to find the learning rate (6e-5), the number of LSTM layers (4), and the number of units (128) of the *body model* (498,409 trainable parameters). We used the same learning rate and number of units for the *arm model*, and performed a grid search to find the number of LSTM layers (5; 607,256 parameters). We used the same learning rate when training the linear models (5,040 and 576 parameters for the *body* and *arm models*, respectively). For the transformer models, we used a positional embedding layer followed by an encoder stack consisting of identical layers with a multi-head self-attention sub-layer and a feed-forward sub-layer [42]. We set the embedding dimension (256), internal dimension of the feed-forward network (1024), and attention and value key size (64) based on previous work with movement data [43], and performed a grid search to find the number of layers (2) in the encoder stack and the number of self-attention heads (4). We used a custom learning rate scheduler according to [42] and the same hyper-parameters for both *arm* (1,591,832 parameters) and *body* (1,618,793 parameters) *models*. For all three model architectures, we used a batch size of 64, the Adam optimizer [44], and a weighted mean squared error as loss function. We observed improved kinematic accuracy by doubling the weight on the error associated with the three markers located on each foot, while keeping the weight for all other markers at one. This can be explained by the sensitivity of the ankle and subtalar angles to small changes in foot marker positions. All results presented here are based on this loss function, with doubled weights for the foot markers.

For each dataset, we divided the data into three subsets: training (80%), validation (10%), and test (10%) sets. This partitioning was conducted on a per-subject basis to ensure that data from a subject was exclusively assigned to one set. We used an early stopping criterion (delta of 0 and patience of 3) based on the validation loss and fine-tuned hyperparameters to minimize the root mean squared error (RMSE) on the validation set. Each model was trained twice, and we selected the one with the lowest RMSE on the validation set. Finally, we assessed the performance of the chosen models on the test set using RMSE of non-normalized data. We trained the models in Python 3.11, using Tensorflow 2.12, and one NVIDIA GeForce RTX 3090 GPU.

### C. Performance evaluation

We evaluated the performance of the three marker enhancers through two tasks. First, we assessed their accuracy when computing joint kinematics from videos on a set of benchmark movements. Second, we assessed their ability to generalize across a broad range of movements. We refer to these tasks as *accuracy* and *generalizability* tasks, respectively.

#### 1) Accuracy task

We incorporated the marker enhancers as part of OpenCap to estimate joint kinematics from videos. The OpenCap pipeline consists of four main steps: 1) pose estimation to identify 2D keypoints from videos, 2) triangulation to reconstruct the 3D position of the keypoints, 3) marker enhancement to predict the 3D position of the anatomical markers, and 4) scaling and inverse kinematics in OpenSim to compute joint kinematics from anatomical marker positions. We compared video-(i.e., OpenCap-) based joint kinematics against reference values from a traditional marker-based pipeline in OpenSim using the OpenCap dataset [1], which was not used to train the marker enhancers. The dataset contains synchronized videos, marker-based motion capture marker trajectories, and force plate data from 10 subjects performing four types of activities: walking, squat, sit-to-stand, and drop jump. Each activity is performed in a self-selected manner and in a modified way to simulate the effect of musculoskeletal conditions (e.g., walking naturally and with a trunk sway gait modification). In total, the dataset includes 160 trials (six walking trials, two times five squats and five sit-to-stands, and six drop jumps per subject). We computed joint kinematics from videos using the three marker enhancers (linear, LSTM, and transformer), along with the marker enhancer from [1]. Additionally, we evaluated the direct use of keypoints as input to scaling and inverse kinematics in OpenSim, bypassing marker enhancement, to assess its significance in the pipeline. Both video- and marker-based pipelines employed the same musculoskeletal model [1], [29], [45], [46], comprising 33 degrees of freedom. Our analysis focuses on 21 lumbar and lower-body degrees of freedom (lumbar [3], pelvis in the ground frame [6], hips [2×3], knees [2×1], and ankles [2×2]). For the video-based pipeline, we utilized HRNet (person model: `faster_rcnn_r50_fpn_coco, pose model: hrnet_w48_coco_wholebody_384×288_dark_plus`) from mmpose [47] as the pose estimation model and videos captured from two iPhones (12 Pro; Apple Inc., Cupertino, CA, USA) positioned at approximately ± 45° from the subject’s forward-facing direction. We evaluated video-against marker-based pipelines using RMSE of joint kinematics.

#### 2) Generalizability task

We used marker-based motion capture data from the Total Capture dataset [48], which was not used to train the marker enhancers, to create synthetic video keypoints and evaluate the generalizability of the marker enhancers across a range of movements. We incorporated data from five subjects engaged in movements including performing various ranges of motion, walking freely within a capture volume (i.e., not only walking straight along the world frame’s forward direction), acting, and engaging in freestyle movements. The latter especially covered a broad range of movements (e.g., rolling and doing push ups), on which OpenCap was known to perform poorly. In total, we included data from 38 trials, corresponding to about 0.6 hours of movement. We used a similar approach as for the marker enhancer dataset to generate corresponding synthetic video keypoints and anatomical markers. First, we processed the marker data from motion capture with AddBiomechanics to obtain scaled OpenSim models and coordinate files. We then attached virtual markers corresponding to the video keypoints and (reference) anatomical markers (Fig. 1) to the scaled models and extracted the 3D trajectory of each virtual marker for each coordinate file. Next, we added Gaussian noise (standard deviation: 18 mm) to each time step of the keypoint positions [17]–[19] and passed these positions as input to the marker enhancers to predict anatomical marker positions. Finally, we computed joint kinematics from anatomical marker positions using scaling and inverse kinematics in OpenSim. We used RMSE to compare joint kinematics computed from the reference anatomical markers and from the predicted anatomical markers. We also evaluated the direct use of video keypoints, with and without noise, as input to scaling and inverse kinematics in OpenSim to compute joint kinematics.

### D. Muscle-driven tracking simulations

We evaluated the effect of improved kinematics on the accuracy of dynamic measures (i.e., ground reaction forces and joint moments). To compute dynamics, we generated tracking simulations of joint kinematics using the musculoskeletal model [1], [49]–[51]. The model is driven by 80 muscles actuating the lower-limb joints and 13 ideal torque motors actuating the lumbar, shoulder, and elbow joints. External forces are modeled through six foot-ground contact spheres attached to the foot segments. We formulated the simulations as optimal control problems that aim to identify muscle excitations that minimize a cost function subject to constraints describing muscle and skeleton dynamics. The cost function J includes effort terms (squared muscle activations *a* and excitations of the ideal torque motors *e*_*T*_) and kinematic tracking terms (squared difference between simulated and experimental data), namely tracking of experimental joint positions 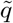, velocities 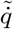, and accelerations 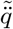:

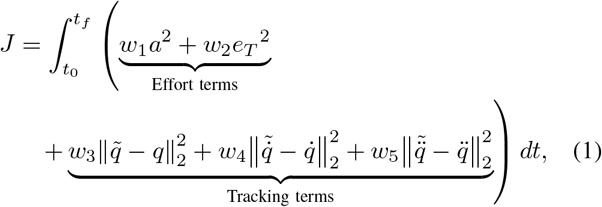

where *t*_0_ and *t*_*f*_ are initial and final times, *w*_*i*_ with *i* = 1, …, 5 are weights, and *t* is time. More details about the problem formulation can be found in [1]. We formulated the problems in Python (v3.9) with CasADi [52] (v3.5.5), using direct collocation and implicit dynamics [49]. We used algorithmic differentiation to compute derivatives [50] and IPOPT to solve the resulting nonlinear programming problems [53].

We generated tracking simulations of joint kinematics using data from the *accuracy* task (i.e., four movements from the OpenCap dataset). We filtered the kinematic data (walking: 6 Hz, squat: 4 Hz, sit-to-stand: 4 Hz, and drop jump: 30 Hz) and manually tuned the cost term weights following a heuristic process. We further tailored each problem formulation to the movement of interest. The walking simulations were from right heel strike to left toe-off, incorporating time buffers at both the beginning and end of the simulations. These buffers were disregarded during data analysis, but they enhanced the simulation results within the intended time period by providing contextual boundaries. Without buffers, we observed instability at the beginning and end of the simulations. For the simulations of squat, sit-to-stand, and drop jump, which involve large hip and knee flexion, we excluded the contribution of passive muscle forces and added reserve actuators to the hip rotation degree of freedom. We found this to be necessary to prevent non-physiological muscle force contribution in deep flexion. For the sit-to-stand simulations, we added a cost term penalizing the model from lifting its heels, thereby incorporating knowledge of the task-specific human objective into the problem formulation. We also added time buffers, following the same rationale as for walking. For the squat simulations, we segmented the repetitions based on the vertical pelvis position, and imposed periodic pelvis position and speed (i.e., same position and speed at the beginning and end of the repetition). Periodic constraints provide contextual boundaries, and we therefore did not add time buffers. Finally, the drop jump simulations were conducted from 0.3 seconds prior to landing to 0.3 seconds after take-off, encompassing a time buffer around the contact phase, which was the period of focus.

To assess the impact of kinematic accuracy on simulated dynamic measures, we conducted simulations tracking joint kinematics computed from marker data derived through three different methods: 1) predicted from videos using the marker enhancer from [1], 2) predicted from videos using the best performing marker enhancer (LSTM) trained on the enhanced dataset, and 3) measured with marker-based motion capture in the laboratory. Comparing the first two cases allows evaluating the influence of the marker enhancer model on simulated dynamic measures. Comparing simulations tracking video-versus laboratory marker-based joint kinematics helps gauge the potential gains in accuracy by refining kinematic estimates from videos (assuming laboratory marker-based kinematics are the gold standard). This analysis also aids in identifying potential limitations of muscle-driven tracking simulations for estimating dynamic measures. For all 10 subjects, we generated three simulations for each movement type (three walking trials, three squats, three sit-to-stands, and three drop jumps) for each variant (self-selected and modified). The laboratory-based marker data from walking lacked a time buffer following left toe-off, thereby not allowing us to apply the same optimal control formulation as for simulations tracking video-based kinematics. We therefore excluded walking simulations tracking laboratory marker-based kinematics from the analysis. In total, we generated 660 simulations (240 simulations for each video-based case and 180 for the laboratory marker-based case). We used RMSE of ground reaction forces (expressed in percent bodyweight, %BW) and joint moments (expressed in percent bodyweight times height, %BW*ht) as performance metrics, with force plate data and joint moments from laboratory-based inverse dynamics as reference. Note that inverse dynamics results include non-physical pelvis residual forces and moments, whereas muscle-driven simulations are dynamically consistent. Differences between joint moments from inverse dynamics and dynamic simulations are therefore not entirely attributable to errors in the simulation pipeline.

## III. Results

## A. Marker enhancer model training

The transformer model achieved the lowest RMSEs on the test set (*body model*: 8.6 mm; *arm model*: 15.3 mm), marginally outperforming the LSTM model (*body model*: 10.0 mm; *arm model*: 16.3 mm). The linear model performed worst, with RMSEs about twice as large as those of the transformer model (*body model*: 16.5 mm; *arm model*: 31.3 mm).

### B. Performance evaluation

#### 1) Accuracy task

The LSTM and transformer models performed best for rotational degrees of freedom (mean RMSEs: 4.1 ± 0.3° and 4.4 ± 0.4°, respectively) (Table II). The linear and LSTM models performed best for pelvis translations (mean RMSEs: 12.4 ± 1.1 mm and 12.8 ± 1.8 mm, respectively). All three models (linear, LSTM, transformer) trained on the enhanced dataset outperformed mean results obtained with the marker enhancer from [1]. Estimating kinematics from keypoints directly resulted in higher mean RMSEs (9.6 ± 1.5° and 24.6 ± 1.8 mm), with RMSEs for some degrees of freedom as large as 43.1° (lumbar extension) and 45.0 mm (pelvis anterior-posterior translation).

**Table II.**
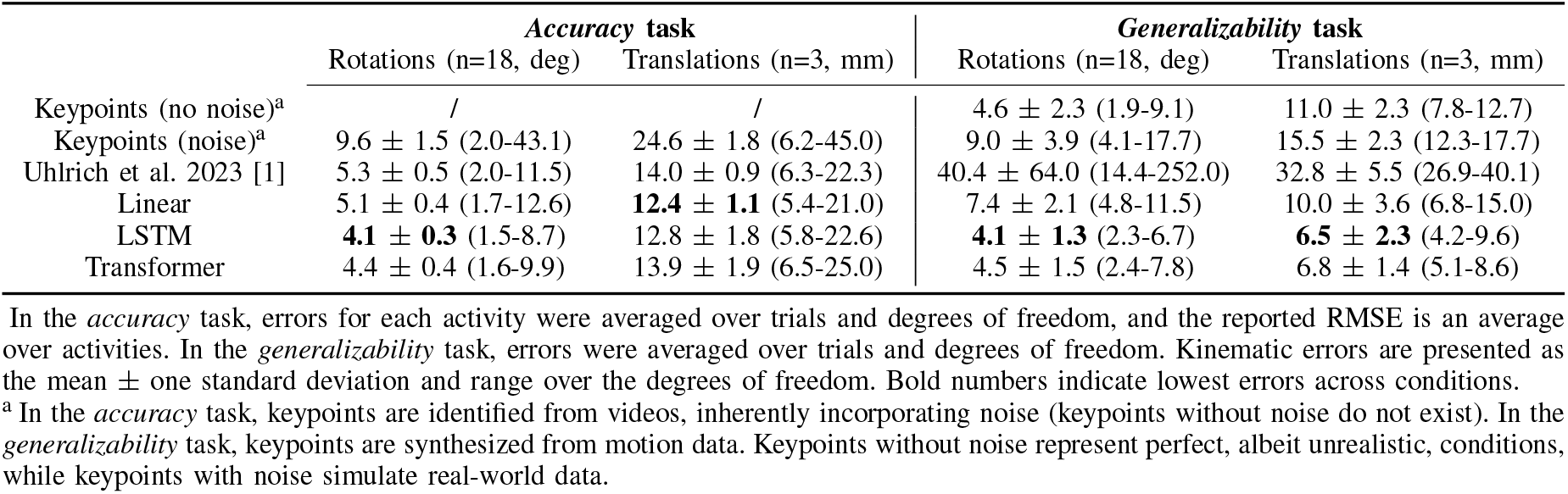
Root Mean Squared Error (rmse) Between Reference and Estimated Joint Kinematics.

#### 2) Generalizability task

The LSTM model achieved the lowest mean RMSEs on the *generalizability* task (Table II) for both rotational degrees of freedom (4.1 ± 1.3°) and pelvis translations (6.5 ± 2.3 mm), marginally outperforming the transformer model (4.5 ± 1.5° and 6.8 ± 1.4 mm). All three models (linear, LSTM, transformer) trained on the enhanced dataset outperformed results obtained with the marker enhancer from [1]. The latter achieved mean RMSEs larger than 40° (mean across rotational degrees of freedom) and 32 mm (mean across pelvis translations), and often failed to capture the movement kinematics (see examples in Fig. 2, column Uhlrich et al. (2023)). All three models also outperformed results obtained when using noisy keypoints instead of anatomical markers as input for scaling and inverse kinematics in OpenSim. When excluding noise, simulating perfect, albeit unrealistic, conditions, the RMSEs decreased but were still larger than those obtained with the LSTM and transformer models.

Overall, the LSTM and tranformer models performed best on both *accuracy* and *generalizability* tasks. We incorporated the LSTM, which is easier to use across different operating systems, into OpenCap. For estimating dynamics using muscle-driven tracking simulations, we also used the kinematics produced with the LSTM as tracking data.

### C. Muscle-driven tracking simulations

On average across movements, the muscle-driven simulations achieved slightly lower RMSEs when tracking the kinematics produced with the LSTM (ground reaction forces: 6.7 ± 4.3 %BW; joint moments: 1.34 ± 0.96 %BW*ht) compared to the kinematics produced with the marker enhancer from [1] (ground reaction forces: 7.2 ± 4.8 %BW; joint moments: 1.37 ± 1.04 %BW*ht) (Table III). Despite some improvements, the changes were limited and not consistent across movements and dynamic quantities. RMSEs consistently showed lower values for drop jumps and higher values for walking, albeit with relatively small changes. Mixed results were observed for squat and sit-to-stand.

**Table III.**
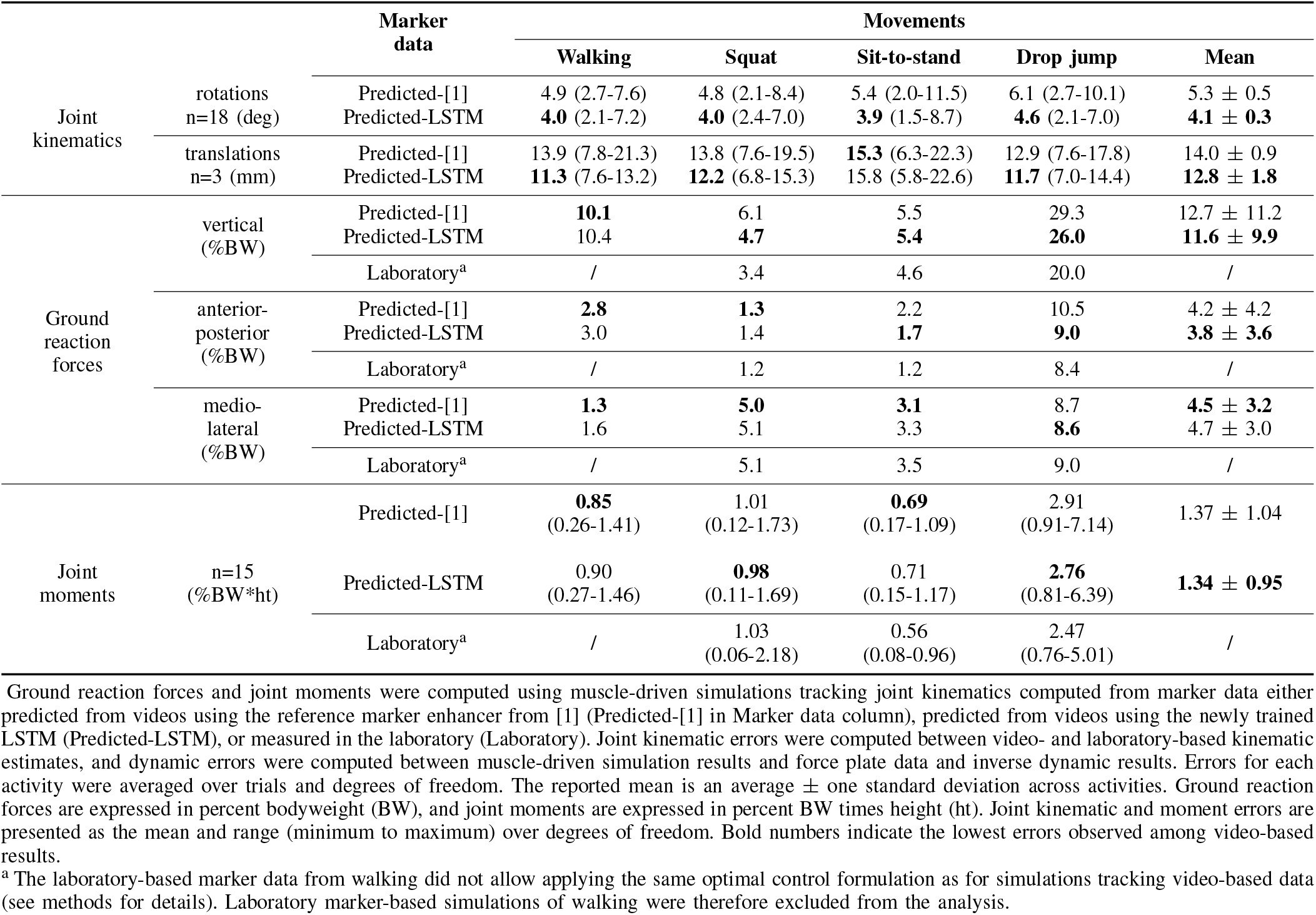
Root Mean Squared Error (RMSE) Between Reference and Estimated Kinematics and Dynamics.

Tracking laboratory-based kinematics did not dramatically and consistently improve the accuracy of dynamic estimates. When tracking kinematics computed from laboratory-instead of video-based marker data, the muscle-driven simulations achieved lower vertical and anterior-posterior ground reaction force RMSEs, but higher medio-lateral RMSEs for squat, sit-to-stand, and drop jumps. This resulted in higher joint moment RMSEs for squat, and lower RMSEs for sit-to-stand and drop jumps.

Out of 660 simulations, seven did not converge and were excluded from the analysis. All other simulations converged to an optimal solution, and we used the same settings for all simulations of a same movement type (e.g., squat).

## IV. Discussion

We created a diverse dataset that enabled the creation of a marker enhancer capable of accurately capturing joint kinematics for a broad range of movements. We demonstrated this generalizability using an unseen dataset comprising a wide array of movements. Furthermore, we showed that the marker enhancer increases accuracy on a benchmark task of estimating joint kinematics from videos. We integrated the marker enhancer in OpenCap, an open-source software platform used by thousands of researchers to measure movement from smartphone videos. Our marker enhancer improves OpenCap’s performance, particularly in tasks where it previously exhibited low accuracy, like when a subject is prone or supine, or changes direction.

We compared different model architectures and found that both LSTM and transformer models achieved similar performance levels on our evaluation tasks, surpassing the performance of a linear model. We hypothesize that the task— predicting a dense set of 3D marker coordinates from a sparse one—is relatively straightforward, allowing both models to achieve comparable results. Both LSTM models, through their memory cells and gated architecture, and transformer models, through their attention mechanisms, leverage the temporal dynamics of time-series data. This utilization of temporal information and the inherent ability of these models to capture non-linear relationships likely contribute to their superior performance compared to a simpler linear model.

The marker enhancer is both accurate and generalizable, and we hypothesize that further accuracy improvements will be challenging to realize without compromising generalizability. We base this assertion on serveral observations. First, our model performs equally well (RMSE of about four degrees) on both the *accuracy* task, which comprises a limited set of controlled movements from the training set, and the *generalizability* task, which comprises a broader range of movements not explicitly covered in training. This consistency suggests that our model is generalizable and not overfitting to specific movements. If it were, we would have expected worse performance on the *generalizability* task compared to the *accuracy* task. Second, our model surpasses the performance of our previous model [1] on the *accuracy* task, despite the fact that our previous model was trained on a smaller and less diverse dataset that comprised the movements tested in this task. This suggests that our new model has not sacrificed accuracy on specific movements for increased generalizability. Otherwise, we would have anticipated a decrease in accuracy on the *accuracy* task compared to our previous model [1]. Third, both the LSTM and transformer models demonstrated similar performance across both evaluation tasks. This outcome suggests that the specific architecture may not be a critical factor in this context, as both models achieved comparable results despite their structural differences. While further accuracy gains can be expected on a given movement (e.g., walking) by fine-tuning the model with a movement-specific dataset, we do not expect that refining the model architecture will lead to substantial accuracy gains on our evaluation tasks.

Our results emphasize the importance of using a dense set of markers to compute joint kinematics. Using sparse keypoints resulted in a larger mean kinematic error (9.6 ± 1.5°) and in a wider range of errors (up to 43.1°) across degrees of freedom (Table II, *Accuracy* task) compared to using dense anatomical markers. Our results on the *generalizability* task (Table II, *Generalizability* task) confirm this claim and further highlight that even under ideal conditions—where we have knowledge of the exact position of the sparse keypoints relative to the multi-body model, thereby ignoring noise inherent to pose estimation models—using sparse keypoints does not result in higher accuracy compared to using the dense set of anatomical markers predicted by the marker enhancer. Interestingly, when using sparse keypoints with noise, thereby simulating more realistic conditions, the error increased and was similar in magnitude (nine degrees) to results on the *accuracy* task with video data. This validates our choice of noise magnitude (standard deviation: 18 mm).

Combining 2D pose estimation models with the marker enhancer is one method for obtaining a dense markerset from videos; however, other approaches are viable. Sárándi et al. [54] introduced a method that integrates diverse datasets used to train pose estimation models, accommodating variations in labeled keypoints—some datasets containing sparser or denser markersets. This approach shows potential for predicting a denser set of keypoints from a sparser one, and is worthy of comparison with our marker enhancer in future studies. Another approach is direct estimation of a dense set of markers from videos, potentially mitigating bias and error accumulation associated with our two-step process (pose estimation followed by marker enhancement). Our preliminary research in this direction [55] involved creating a synthetic video dataset based on the SMPL model [56] with known anatomical marker positions. Using this dataset, we then retrained 2D pose estimation models to directly identify these markers from videos, yielding promising results on the OpenCap dataset used in our *accuracy* task. However, overall performance was not as high as that achieved with our marker enhancer. Future work should concentrate on refining the synthetic video dataset to enhance kinematic accuracy.

Our muscle-driven simulation results indicate that tracking more accurate kinematics does not always result in more accurate dynamics, highlighting the need for better methods to estimate forces. We found that improvements in joint kinematic accuracy obtained with the marker enhancer did not consistently lead to improved estimates of dynamic quantities across the different movements (Table III, Predicted-LSTM versus Predicted-[1]). And even when our muscle-driven simulations tracked joint kinematics computed from gold-standard, laboratory-based marker data (Table III, Laboratory) the dynamics errors, with respect to force plate data and inverse dynamics results, were comparable to those generated when tracking kinematics from video-based data.

We suggest that three main factors contribute to these outcomes: musculoskeletal modeling assumptions, optimal control problem formulation assumptions, and simulation sensitivity. First, modeling assumptions limit the ability of the musculoskeletal model to track given kinematics with physio-logically realistic forces. This causes either the kinematics in the dynamic simulation to deviate from the reference kinematics, or the produced forces to be non realistic. For instance, we found the gluteus maximus muscles to produce excessive hip abduction and external rotation torques in deep hip flexion in activities like squat or sit-to-stand. Personalizing the models, in particular better characterizing the muscle-tendon parameters of the Hill-type muscle model (e.g., using electromyography [22], [57]) and the muscle geometries (e.g., using medical imaging [58]), might improve the muscle operating ranges and force production capabilities, resulting in more realistic force estimates for the given kinematics. Second, the formulation of the optimal control problems underlying the muscle-driven simulations might not fully capture the task constraints and the subject’s motor control objective. For instance, the intention of the subject, which is modeled through the cost function, for walking likely differs from squatting. Using experimental force data with methods like inverse optimal control [59] might help optimize the cost function. It will, however, remain difficult to find a formulation that is robust against how different subjects perform a given task. Third, dynamic simulations lack robustness, which can lead to varying output quality. For example, performing simulations of walking for different trials using the same problem formulation may result in different quality outcomes. While this might relate to different underlying control strategy, we also found the simulations to produce different solutions when, for instance, slightly adjusting the time window of the problem, underlining the simulations’ overall sensitivity. In this analysis, we applied a consistent scaling approach to obtain the musculoskeletal model and used the same problem formulation for all movements of a specific type (e.g., walking) across all ten subjects. Refining the model and problem formulation for each trial could potentially enhance result accuracy. However, this task is challenging without experimental force data available for comparison.

Overall, our muscle-driven tracking simulations provide ground reaction force and joint moment estimates that we previously showed were good enough for applications such as screening for disease risk and informing rehabilitation decisions [1]. Our findings suggest that to improve accuracy, the focus should shift to the methods used for obtaining dynamics rather than enhancing joint kinematic estimates from videos. For example, using data-driven models to predict dynamic quantities like ground reaction forces and inform muscle-driven simulations could be a promising approach.

## V. Conclusion

We created a large and diverse dataset and developed a marker enhancer that is accurate and generalizable across a broad range of movements. Our results illustrated that to improve the accuracy of dynamic estimates from videos using muscle-driven tracking simulations, improving the fidelity of the musculoskeletal model and refining the optimal control problem formulation might have a bigger impact than improving the accuracy of kinematic estimates. By integrating our marker enhancer into OpenCap, we enable its thousands of users to more accurately measure a wider variety of movements from smartphone videos.

## Appendix a Code and Data Availability

Source code, trained models, and public data will be available at https://github.com/antoinefalisse/marker-enhancer.

